# Optical activation of TrkB receptors

**DOI:** 10.1101/2019.12.15.876722

**Authors:** Peiyuan Huang, Aofei Liu, Yutong Song, Jen M. Hope, Bianxiao Cui, Liting Duan

## Abstract

Brain-derived neurotrophic factor (BDNF), via activation of tropomyosin receptor kinase B (TrkB), plays a critical role in neuronal proliferation, differentiation, survival, and death. Dysregulation of TrkB signaling is implicated in neurodegenerative disorders and cancers. Precise activation of TrkB receptors with spatial and temporal resolution is greatly desired to study the dynamic nature of TrkB signaling and its role in related diseases. Here we develop different optogenetic approaches that use light to activate TrkB receptors. Utilizing the photosensitive protein *Arabidopsis thaliana* cryptochrome 2 (CRY2), the light-inducible homo-interaction of the intracellular domain of TrkB (iTrkB) in the cytosol or on the plasma membrane is able to induce the activation of downstream MAPK/ERK and PI3K/Akt signaling as well as the neurite outgrowth of PC12 cells. Moreover, we prove that such strategies are generalizable to other optical homo-dimerizers by demonstrating the optical TrkB activation based on the light-oxygen-voltage domain of aureochrome 1 from *Vaucheria frigida*. The results open up new possibilities of many other optical platforms to activate TrkB receptors to fulfill customized needs. By comparing all the different strategies, we find that the CRY2-integrated approach to achieve light-induced cell membrane recruitment and homo-interaction of iTrkB is most efficient in activating TrkB receptors. The optogenetic strategies presented are promising tools to investigate BDNF/TrkB signaling with tight spatial and temporal control.

## Introduction

Tropomyosin receptor kinase B (TrkB), a preferential receptor for brain-derived neurotrophic factor (BDNF), is expressed in both central and peripheral nervous system[1–3]. Upon binding to BDNF, TrkB receptor undergoes homo-dimerization which consequently leads to the phosphorylation of its cytoplasmic tyrosine kinase domain and activate TrkB signaling[1,4]. BDNF/TrkB signaling is critical for the survival, differentiation, proliferation and plasticity of neuronal cells[5], while defective BDNF/TrkB signaling is implicated in various diseases including neuroblastoma[6], glaucoma[7], Alzheimer’s and Parkinson’s diseases[8,9] and psychiatric depression[10]. The ability to precisely modulate BDNF/TrkB pathway can greatly facilitate the investigation of the role of TrkB signaling in crucial cell functions as well as the pathogenesis of many diseases.

Optogenetics emerges as a powerful tool that employs light to control cellular activities. Optogenetic modules enabling light-inducible protein-protein interactions have been widely used to control a variety of signaling pathways[11,12], including the mitogen-activated protein kinase/extracellular signal-regulated kinase (MAPK/ERK) pathway[13,14], phosphatidylinositol 3-kinase/protein kinase B (PI3K/Akt) pathway[15,16], TGF-β signaling[17], receptor tyrosine kinases (RTKs) signaling[18], and G protein-coupled receptors signaling[19,20]. The use of light permits unprecedented spatiotemporal precision to modulate signaling events, compared to conventional methods such as pharmacological perturbation. There have been several reports that exploits light to control TrkB receptors by optically inducing the homo-dimerization of full-length TrkB receptors or membrane-bound intracellular domains of TrkB receptors[21–23]. However, the full-length receptor fused to photosensory proteins is still responsive to ligands via its extracellular domain. In addition, whether the membrane-bound light-responsive TrkB possesses the best optical activation efficacy is unclear, as we previously found that light-induced membrane recruitment and homo-interaction of the intracellular domain of tropomyosin receptor kinase A (iTrkA) is more effective to activate TrkA signaling than the optical homo-interaction of membrane-bound iTrkA[24]. Here in this report, we develop and compare different optical approaches to regulate TrkB signaling (OptoTrkB) and inspect their efficiency. Based on the light-gated homo-association of *Arabidopsis thaliana* cryptochrome 2 (CRY2) and hetero-dimerization of CRY2 and the truncated version of basic helix-loop-helix protein *Arabidopsis* CIB1 (CIBN)[25], we show that photo-mediated homo-interaction of TrkB intracellular domain (iTrkB), either in the cytosol or on the cell membrane, can activate the downstream signaling cascades of TrkB. In addition, we prove the light-induced homo-interaction of iTrkB is a general method to activate TrkB receptor that can be combined with other photosensory proteins. We demonstrate that the homo-binding of iTrkB actuated by the light-oxygen-voltage domain of aureochrome 1 from *Vaucheria frigida* (AuLOV) is also able to trigger TrkB signaling. Furthermore, we compare all the optogenetic strategies and determine that the light-induced relocation and homo-association of iTrkB on the cell membrane utilizing CRY2/CIBN system is most efficient in activating TrkB downstream signaling and PC12 neurite growth.

## Results

### Design of OptoTrkB systems using CRY2/CIBN modules

Endogenous TrkB receptor consists of an extracellular domain working as the binding site of neurotrophin, a transmembrane domain, and an intracellular catalytic domain[1]. In this study, we use the truncated intracellular domain of TrkB (a.a. 455-822, iTrkB) with the extracellular and transmembrane domain removed, so that the engineered receptors will be inert to TrkB ligands. The CRY2 used in this study is the photolyase homology (PHR) domain (a.a. 1-498), and CIBN is the truncated version of CIB1 (a.a. 1-170)[26].

In the first strategy named as Opto-iTrkB (Fig. 1A), iTrkB is fused to the photosensitive protein CRY2 and is distributed in the cytoplasm. In the second strategy denoted as Lyn-Opto-iTrkB (Fig. 1B), with an N-terminal Lyn tag working as a membrane-targeting peptide, the fusion protein is localized on the plasma membrane. The third strategy, named as Opto-iTrkB + CIBN-CAAX, contains two components, Opto-iTrkB and CIBN fused to CAAX (Fig. 1C), where the CAAX motif targets to the cell plasma membrane[27]. Blue light can recruit Opto-iTrkB to plasma membrane via CRY2-CIBN association, and then induce the interaction of Opto-iTrkB via CRY2-CRY2 binding. In the three strategies, light-induced CRY2 homo-interaction is expected to drive the association of iTrkB in cell cytoplasm or on the cell membrane, which consequently triggers the autophosphorylation of iTrkB and the activation of TrkB receptors.

**Fig. 1.**
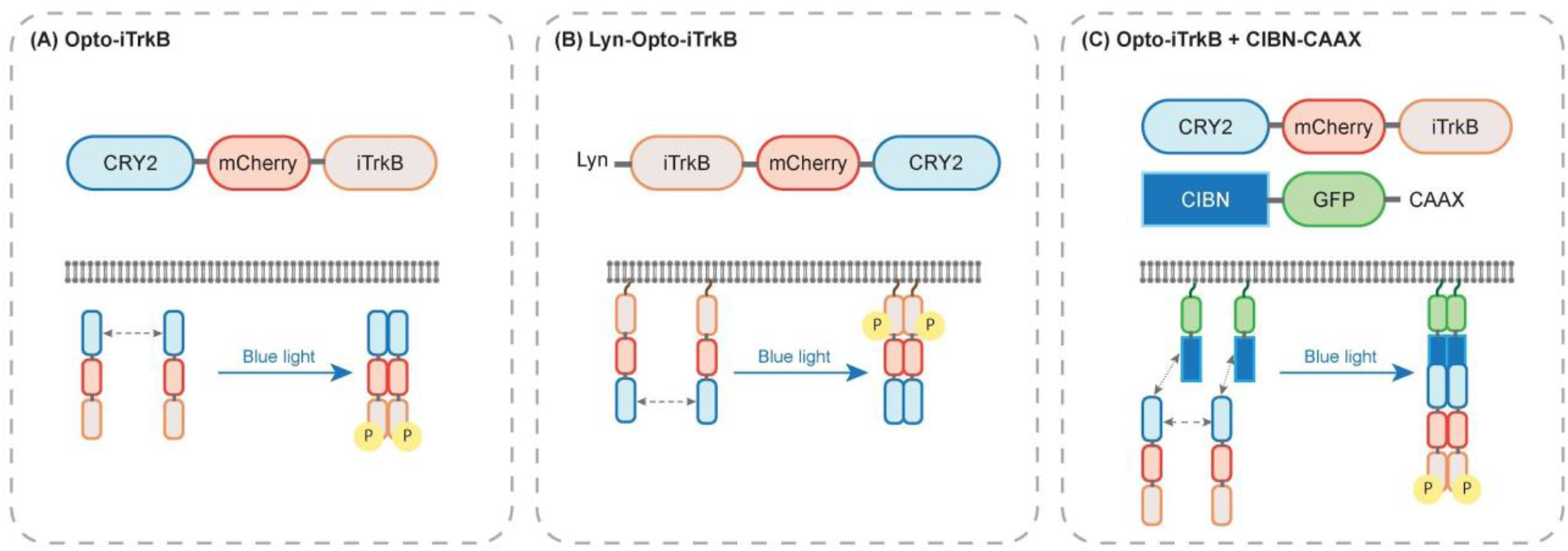
Design of three OptoTrkB systems based on light-induced CRY2-CRY2 homo-interaction and CRY2-CIBN hetero-dimerization. (A) Opto-iTrkB: Photosensitive protein CRY2 is fused to the N-terminus of the intracellular domain of TrkB (iTrkB). (B) Lyn-Opto-iTrkB: An N-terminal Lyn membrane-targeting sequence is linked to the N-terminus of iTrkB while CRY2 is appended to the C-terminus of iTrkB. (C) Opto-iTrkB + CIBN-CAAX: CIBN-CAAX is localized on the plasma membrane, which recruits Opto-iTrkB to the cell membrane upon blue light exposure.

### OptoTrkB systems activate MAPK/ERK and PI3K/Akt pathways

To evaluate the OptoTrkB systems, we first confirmed the expression and distribution of the optogenetic fusion proteins. We inserted a red fluorescent protein mCherry (mCh)to visualize the distribution of OptoTrkB fusion proteins. In NIH 3T3 cells expressing Lyn-iTrkB-mCh-CRY2, the fusion protein Lyn-Opto-iTrkB was mostly localized on the plasma membrane (Fig. S1A). We also confirmed that in cells expressing Opto-iTrkB + CIBN-CAAX, CRY2-mCh-iTrkB was quickly recruited to plasma membrane upon blue light illumination, validating the photo-mediated CRY2-CIBN dimerization (Fig. S1B).

Next, we set out to assay the light-induced activation of TrkB signaling cascades. TrkB activation triggers downstream signaling cascades including MAPK/ERK pathway and PI3K/Akt pathway. We employed the nucleus export of kinase translocation reporter (KTR) to report the activation of MAPK/ERK pathway. The KTR technology was developed to report phosphorylation-regulated nucleocytoplasmic translocation[28], where the relative intensity of cytoplasmic versus nuclear fluorescence is used as a proxy for kinase activity in living cells[29]. On the other hand, subcellular translocation of the pleckstrin homology domain of Akt (AktPH) was used to indicate the activation of PI3K/Akt pathway, where AktPH-GFP can be recruited to the cell membrane upon the phosphorylation of Akt[30]. NIH 3T3 cells were transfected with each of the three OptoTrkB systems together with ERK-KTR-GFP or AktPH-GFP, respectively. Blue light pulses at 5-s intervals (470 nm, 200 ms pulse duration at 9.7 W/cm^2^) was used to both visualize the redistribution of ERK-KTR-GFP or AktPH-GFP and stimulate the OptoTrkB systems. Upon the delivery of blue light, obvious translocation of ERK-KTR-GFP from the cell nucleus to the cytoplasm was observed in all the three systems with the GFP signal significantly decreased inside the cell nucleus, which confirms the activation of MAPK/ERK signaling (Fig. 2A). Moreover, in all the three OptoTrkB systems, AktPH-GFP was successfully recruited to plasma membrane under blue light illumination with the GFP intensity on the plasma membrane increased, indicating the activation of PI3K/Akt signaling (Fig. 2B). As a control, there was no noticeable translocation of either ERK-KTR-GFP or AktPH-GFP observed in cells only expressing CIBN-CAAX, which proves that light-actuated homo-interaction of iTrkB indeed triggers the downstream signaling cascades (Fig. S2).

**Fig. 2.**
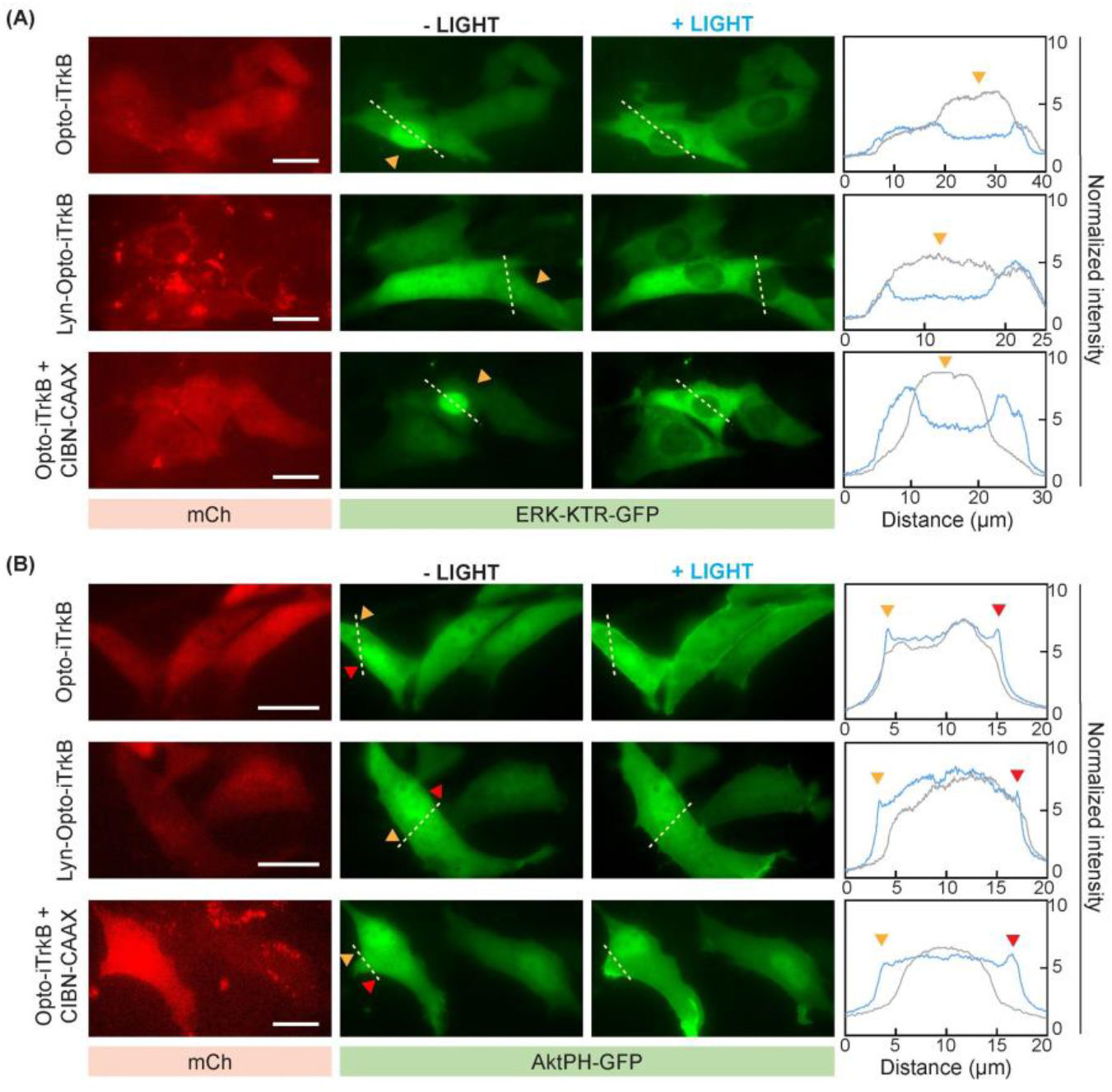
OptoTrkB systems activate the downstream MAPK/ERK and PI3K/Akt pathways. Blue light pulses (200 ms, 9.7 W/cm^2^) were delivered at 5 s intervals for 10 min. (A) Opto-iTrkB, Lyn-Opto-iTrkB and Opto-iTrkB + CIBN-CAAX activates MAPK/ERK pathway, assayed by ERK-KTR-GFP translocating from nucleus to cytosol. Right panels show the normalized intensity change along dashed lines before (black line plot) and after blue light stimulation (blue line plot). (B) Opto-iTrkB, Lyn-Opto-iTrkB and Opto-iTrkB + CIBN-CAAX activates PI3K/Akt pathway, assayed by membrane translocation of AktPH-GFP. Right panels show the normalized intensity change along dashed lines before (black line plot) and after blue light stimulation (blue line plot). Scale bars, 20 μm.

### Homo-interactions of iTrkB actuated by another optical homo-dimerizer, AuLOV, can also activate TrkB receptors

After validating the activation of TrkB receptors by CRY2-based OptoTrkB systems, we asked whether the optical approaches can be generalizable to other light-gated homo-dimerizers. While CRY2 is widely used to actuate homo-interactions of target proteins, a number of other optical homo-dimerizers have been developed and applied in various optogenetics applications[31], including *Arabidopsis* URV8[32], *Neurospora crassa* VVD[33,34], *Vaucheria frigida* AuLOV[35], *Erythrobacter litoralis* EL222[36], and cyanobacterial phytochrome CPH1[37]. Inspired by a recent report that investigates the activation of TrkA signaling using AuLOV[38], we hypothesized that the homo-binding of AuLOV can also be leveraged to achieve optical activation of TrkB. We designed two AuLOV-incorporated OptoTrkB strategies (Fig. 3A). In the first strategy, AuLOV was fused to the N-terminus of iTrkB, similar to the CRY2-based Opto-iTrkB. The second strategy fused a membrane-targeting Lyn tag to the N-terminus of iTrkB and AuLOV to the C-terminus of iTrkB, similar to the CRY2 version of Lyn-Opto-iTrkB. Using translocation assays of ERK-KTR-GFP or AktPH-GFP, we confirmed that two AuLOV-based OptoTrkB systems can successfully activate TrkB downstream signaling pathways. NIH 3T3 cells were co-transfected with ERK-KTR-GFP or AktPH-GFP along with either AuLOV-mCh-iTrkB or Lyn-iTrkB-AuLOV-mCh. Under the same blue light stimulation (470 nm pulses at 5-sec intervals for 10 mins, 200 ms pulse duration at 9.7 W/cm^2^), ERK-KTR-GFP successfully redistributed out of the cell nucleus, while AktPH-GFP accumulated more on the cell membrane (Fig. 3B). Therefore, AuLOV-integrated OptoTrKB systems can also effectively activate the downstream signaling of TrkB, which implies that light-actuated iTrkB homo-association is a general method to stimulate TrkB signaling.

**Fig. 3.**
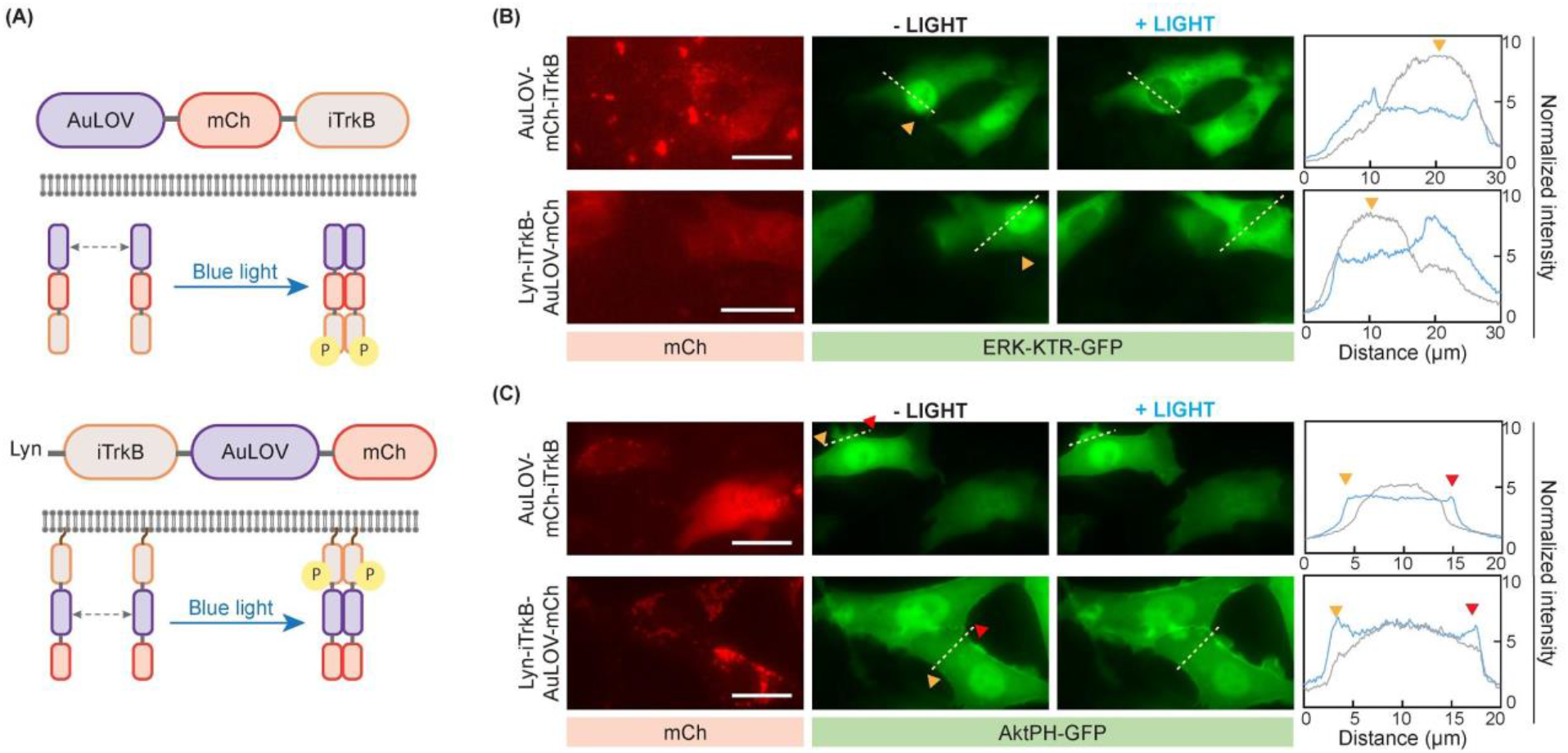
Homo-interactions of iTrkB actuated by AuLOV can also activate TrkB signaling. (A) Illustrations of AuLOV-based OptoTrkB systems. AuLOV is fused to the N-terminus of iTrkB, forming AuLOV-mCh-iTrkB. In the second system, the membrane tag Lyn is attached to the N-terminus of iTrkB and AuLOV is connected to the C-terminus of iTrkB, forming Lyn-iTrkB-AuLOV-mCh. (B) AuLOV-mCh-iTrkB or Lyn-iTrkB-AuLOV-mCh activates MAPK/ERK pathway, indicated by ERK-KTR-GFP translocation from the nucleus to the cytosol. Right panels show the normalized intensity change before (black line plot) and after blue light stimulation (blue line plot). (C) AuLOV-mCh-iTrkB or Lyn-iTrkB-AuLOV-mCh activates PI3K/Akt pathway, shown by AktPH-GFP translocation onto the cell membrane. Right panels show the normalized intensity change before (black line plot) and after blue light stimulation (blue line plot). Scale bars, 20 μm.

### Light-induced OptoTrkB activation promotes neurite growth in PC12 cells

It has been shown that the activation of TrkB signaling with BDNF can elicit PC12 cell neurite growth[39,40]. To examine the capability of our light-activatable OptoTrkB systems in recapitulating the BDNF function, we tested whether light can induce the differentiation of PC12 cells expressing OptoTrkB systems in the absence of neurotrophic factor.

PC12 cells expressing each of CRY2- or AuLOV-based OptoTrkB systems were subject to intermittent 200 μW/cm^2^ blue light (5 sec on, 5 sec off) for 24 hours. CIBN-GFP-CAAX was used to visualize the morphology in cells expressing Opto-iTrkB + CIBN-CAAX system. The membrane-tagged GFP-CAAX was expressed in cells expressing other OptoTrkB systems to mark cell shapes. Cells only transfected with CIBN-GFP-CAAX was used as a control group. Another set of PC12 cells undergoing the same transfection treatment was kept in the dark as a dark control. Varied extents of neurite growth were observed among all the OptoTrkB groups (Fig. 4A-E). As a comparison, cells expressing only CIBN-GFP-CAAX had no obvious neurite outgrowth in the presence of light stimulation (Fig. 4F). In addition, cells expressing each of the OptoTrkB systems that were kept in the dark did not grow noticeable neurites (Fig. S3), which proves that the PC12 cell differentiation is indeed resulted from light-induced TrkB activation.

**Fig. 4.**
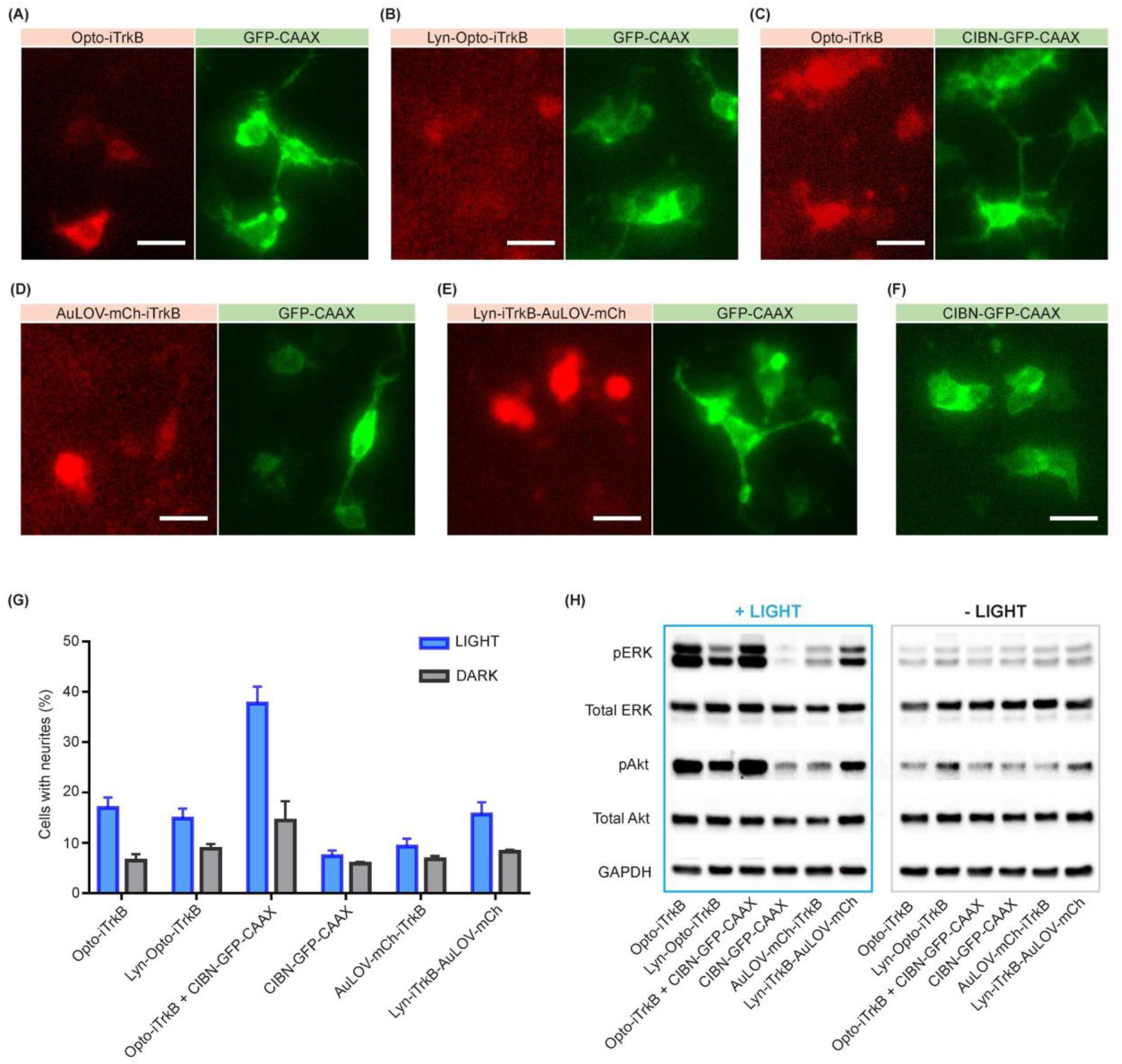
Opto-iTrkB + CIBN-CAAX strategy based on CRY2 shows the best efficiency in activating PC12 neurite growth and downstream signaling pathways. (A-F) OptoTrkB systems promote neurite growth in PC12 cells in the absence of BDNF. In PC12 cells expressing each of the OptoTrkB systems (A-E), noticeable neurite growth was observed in some cells. However, PC12 cells only transfected with CIBN-GFP-CAAX did not show obvious neurite growth. (G) Quantification of the percentage of cells with neurites indicates that the CRY2-based Opto-iTrkB + CIBN-CAAX strategy is most effective to trigger neurite outgrowth. Cells were counted if they grew neurites >10 μm long. Results are averaged from over 300 cells in each group across three independent sets of experiments. (H) Western blot analysis to probe the levels of phosphorylated ERK (pERK) and phosphorylated Akt (pAkt) shows that Opto-iTrkB + CIBN-CAAX can induce the highest levels of pERK and pAkt. Cells were exposed to continuous blue light at 400 μW/cm^2^ for 20 mins or left in the dark. Scale bars, 20 μm.

### Opto-iTrkB + CIBN-CAAX can most efficiently activate TrkB receptors

After verifying the optical activation of TrkB via each OptoTrkB system, we next set out to determine which system can most effectively activate TrkB signaling. We compared the activation efficiency by two means, the extents of light-induced PC12 cell neurite growth and the levels of phosphorylated downstream signaling proteins assayed by western blotting.

First, to quantify the light-induced PC12 cell differentiation, we counted the percentage of cells with neurite growth among all the transfected cells in each experimental group described above (Fig. 4G). Cells with neurites longer than 10 μm were counted and the results are presented as averages from three independent experiments (details available in Supplementary Table S1). In the presence of blue light illumination, only 7% of cells expressing CIBN-GFP-CAAX alone showed noticeable neurites. As a comparison, with light stimulation, cells expressing either CRY2- or AuLOV-based OptoTrkB system had more neurite outgrowth than the CIBN-GFP-CAAX control group. In the same transfection groups kept in the dark, few cells possessed noticeable neurites (Fig. S3). Particularly, in Opto-iTrkB + CIBN-CAAX group, 38% of cells grew neurites upon light exposure, which showed the highest percentage among all the groups. On the other hand, similar portions of cells obtained noticeable neurites in Opto-iTrkB (17%), Lyn-Opto-iTrkB (15%) or Lyn-iTrkB-AuLOV-mCh (16%) group under light stimulation. In cells transfected with AuLOV-mCh-iTrkB, only 9% of cells had neurite growth, which is slightly higher than CIBN-GFP-CAAX control group. We also measured the average numbers of neurite branches in differentiated cells in each light-illuminated group (Fig. S4), which showed no obvious difference among all the groups. In summary, the quantification results indicate that the CRY2-based Opto-iTrkB + CIBN-CAAX is the most effective strategy in inducing PC12 cell differentiation.

Next, we examined the levels of phosphorylated ERK1/2 (pERK, Thr202 and Tyr204) and phosphorylated Akt (pAkt, Ser473) among different groups by immunoblotting (Fig. 4H). After being transfected with each of the OptoTrkB systems and treated with overnight serum-starvation, PC12 cell cultures were subjected to continuous blue light stimulation (400 μW/cm^2^) for 20 minutes, followed by immediate cell lysis. One control group of PC12 cells was only transfected with CIBN-GFP-CAAX. Another set of PC12 cell cultures was kept in the dark to check the dark activation. Lysed samples were probed for total ERK1/2, pERK, total Akt and pAkt, while GAPDH was probed for a loading control (Fig. 4H). In the absence of blue light illumination, minimal levels of pERK and pAkt were detected in all groups, with Lyn-Opto-iTrkB and Lyn-iTrkB-AuLOV-mCh gaining slightly higher dark activation. Contrarily, blue light illumination induced a significant rise of pERK and pAkt levels in the groups of cells expressing Lyn-Opto-iTrkB or Lyn-iTrkB-AuLOV-mCh, an even greater elevation in the Opto-iTrkB or Opto-iTrkB + CIBN-CAAX groups, and a slight increase in the group of AuLOV-mCh-iTrkB. Particularly, cells expressing Opto-iTrkB + CIBN-CAAX obtained the highest light-induced levels of pERK and pAkt among all the groups.

As a negative control, the levels of pERK and pAKt remained low in the light-illuminated CIBN-GFP-CAAX group. Quantification of western blotting results confirms that the Opto-iTrkB + CIBN-CAAX strategy can lead to the strongest activation of TrkB downstream signaling among all OptoTrkB systems (Fig. S5).

## Discussion

In this study, we presented five different optical strategies that activate TrkB receptors through light-induced homo-binding of iTrkB, which utilizes either CRY2/CIBN system or AuLOV. These light-activatable TrkB receptors are insensitive to BDNF as well as other TrkB agonists or antagonists that act via the receptor extracellular domain. We demonstrated that the five optical strategies can activate TrkB downstream signaling cascades and evoke neurite growth of PC12 cells. Particularly, we found that CRY2-based Opto-iTrkB + CIBN-CAAX is the most effective strategy to activate TrkB receptors by quantifying the degrees of PC12 neurite outgrowth and comparing the levels of phosphorylated ERK and Akt. The result agrees with our previous report that the same design for TrkA receptors, Opto-iTrkA + CIBN-CAAX, can activate TrkA signaling better than other methods exploiting full-length receptors or membrane-bound intracellular domain of the receptors[24]. To further improve the activation efficiency in the future, CRY2 variants with enhanced homo-interacting abilities, such as CRY2high[41], CRY2clust[42] or CRY2oligo[43], can be incorporated into the Opto-iTrkB + CIBN-CAAX construct. We expect that the Opto-iTrkB + CIBN-CAAX strategy would be a powerful tool to interrogate TrkB signaling.

Among the five strategies, Lyn-Opto-iTrkB and Lyn-iTrkB-AuLOV-mCh systems share the same design to optically induce the homo-interaction of membrane-bound iTrkB. The two strategies led to similar extents of PC12 neurite growth and phosphorylation of downstream signaling proteins. Though both Opto-iTrkB and AuLOV-mCh-iTrkB can induce the association of iTrkB in the cytosol, Opto-iTrkB excels in the activation of TrkB signaling possibly due to the strong homo-oligomerization capability of cytosolic CRY2. It has been reported that under the right experimental conditions, the tendency of CRY2 to homo-oligomerize is so strong that cytosolic CRY2 can form large clusters[44]. Interestingly, for the activation efficacy, Opto-iTrkB is more potent than Lyn-Opto-iTrkB while Lyn-iTrkB-AuLOV-mCh is stronger than AuLOV-mCh-iTrkB. To explain such contrast, further investigation on how CRY2 or AuLOV behaves differently in cytosol or on the membrane may be helpful.

We also proved that the light-induced homo-interaction of iTrkB either in the cytosol or on the cell membrane is a general way to activate TrkB signaling. Among assorted optical homo-dimerizers, we demonstrated that CRY2 and AuLOV can be applied to activate TrkB receptors. We believe that this general approach can also be actuated by other homo-dimerizers. For example, homo-dimerizers of different kinetics and binding affinities can be chosen to construct customized OptoTrkB systems. In addition, homo-dimerizers responsive to different wavelengths of light, such as CPH1 that is activatable by red light or UVR8 that can be stimulated by UV light, can be used to satisfy distinct illumination conditions. For example, to construct an orthogonal platform to independently regulate TrkB signaling and other cellular process, a homo-dimerizer with excitation spectra well separated from another optogenetic manipulation of cellular event can be chosen and embedded into our OptoTrkB system. Therefore, our results open up possibilities of many other new optoTrkB systems to fit various needs.

## Materials and Methods

### Plasmid cloning

#### Lyn-iTrkB-mCh-CRY2

The intracellular domain of TrkB (a.a. 455-822) was inserted into our previous construct Lyn-iTrkA-mCh-CRY2[24] at SacI and BsrGI sites using In-Fusion (Clontech).

#### CRY2-mCh-iTrkB

iTrkB was inserted into our previous construct CRY2-mCh-iTrkA[24] at BsrGI and MunI sites using In-Fusion (Clontech).

#### Lyn-iTrkB-AuLOV-mCh

iTrkB was inserted into Lyn-TrkAICD-AuLOV-GFP[38], a kind gift from Prof. Kai Zhang at University of Illinois at Urbana-Champaign, at EcoRV and SacI sites using In-Fusion (Clontech).

#### AuLOV-mCh-iTrkB

AuLOV was inserted into pmCherry-N1 (Clontech) at EcoRI and BamHI using In-Fusion (Clontech). iTrkB was then inserted at BsrGI and MunI using In-Fusion (Clontech).

### Cell culture and transfection

NIH 3T3 cells were cultured in DMEM medium (Thermo Fisher Scientific) supplemented with 10% FBS (fetal bovine serum, Clontech) and 1% P/S (Penicillin-Streptomycin, Thermo Fisher Scientific). PC12 cells were maintained in F-12K medium (Thermo Fisher Scientific) supplemented with 15% v/v horse serum (Thermo Fisher Scientific), 2.5% FBS and 1% P/S. All cell cultures were maintained at 37°C with 5% CO_2_. All cells were transfected with desired DNA plasmids using Lipofectamine 2000 (Thermo Fisher Scientific) according to the manufacturer’s protocol. Transfected cells were allowed to recover and express the desired proteins overnight in complete culture medium. For all experiments, NIH 3T3 cells and PC12 cells were kept in the dark in serum free medium for over 8 hours prior to any light illumination.

### Light stimulation for cell culture

Transfected PC12 cells were allowed for overnight recovery to express the protein, followed by 8 hours serum-starvation the dark to minimize interference from growth factors in the serum. Cells were then illuminated with 200 μW/cm^2^ intermittent blue light (5 sec on/5 sec off) for 24 hours using a custom-built LED array inside a CO2 incubator. Similar illumination conditions have been tested in previous reports and shows no obvious toxicity[24,41,45]. Additional sets of transfected cells were kept in the dark as control groups. Cells were washed with cold PBS and fixed with 4% paraformaldehyde prior to imaging under fluorescence microscope.

### Imaging

All imaging was performed on a Leica DMi8 S microscope. NIH 3T3 Cells were plated on 35 mm confocal dishes with hole size of 13Ø or 20Ø (SPL) and allowed to grow overnight before transfection. After desired treatment, cells were imaged at excitation wavelength of 470 nm (GFP) and 510 nm (mCherry) at 5-sec intervals (200 ms pulse duration at 9.7 W/cm^2^). The light intensity was measured by THORLABS PM100D optical power and energy meter right above the 100x objective.

### Immunoblotting

PC12 cells were transfected by electroporation using the Amaxa Nucleofector II (Lonza). Cells were added to a suspension of DNA in electroporation buffer (7 mM ATP, 11.7 mM MgCl2, 86 mM KH2PO4, 13.7 mM NaHCO3, 1.9 mM glucose), transferred to a 2mm electroporation cuvette (Fisher Scientific) and subjected to the manufacturer provided protocol for PC12 cells. Cells were allowed to recover in culture medium for 40h before serum starvation for 8h prior to experiment.

For the western blot, cells were exposed to blue light at 400 μW/cm2 for 10 minutes before being moved to ice, rinsed with cold PBS and lysed in RIPA buffer. (25 mM Tris HCl, 150mM NaCl, 1% Triton X-100, 1% Sodium deoxycholate, 0.1% SDS) supplemented with protease and phosphatase inhibitor cocktails (Roche 04906837001 and 04693132001). Clarified lysates were mixed with Laemmli sample buffer (Bio-Rad 1610747) and β-mercaptoethanol and boiled for 5min. Lysed samples were subjected to electrophoresis using Bio-Rad’s Mini-PROTEAN system (1658026FC). After separation, protein was transferred to a nitrocellulose membrane (Bio-Rad), followed by standard blotting procedure. Primary antibodies were obtained from Cell Signaling Technology: anti-pAKT (T308) (CST 9275), anti-AKT (CST 9272), anti-pERK1/2 (T202+Y204) (CST 9101), anti-ERK1/2 (CST 9102), and anti-GAPDH (CST 2118). HRP-conjugated secondary antibody (CST 7074) was used for protein band detection. Protein bands were visualized by chemiluminescence (Bio Rad 1705060) using a ChemiDoc imaging system.

### Data analysis

ImageJ software (NIH) was used to process and analyze images. The intensity along the dotted line was plotted and normalized to average background intensity in figures of NIH 3T3 cells. To quantify the neurite growth of PC12 cells, all transfected cells and cells bearing neurites longer than 10 μm were manually counted by ImageJ cell counter. Western blotting results were quantified by densitometry analysis using ImageJ.

## Author contribution

LD and PH conceived the project and designed the experiments. PH, AL, JMH and YS performed the experiments, and PH analyzed data. PH and LD wrote the manuscript. All authors contributed to the discussion of the results.

## Acknowledgement

We thank Dr. Kai Zhang (University of Illinois at Urbana-Champaign) for providing plasmid Lyn-TrkAICD-AuLOV-GFP. This work was supported by a Direct Grant from the Chinese University of Hong Kong (4055095), Shun Hing Institute of Advanced Engineering (SHIAE) Grant (4720247) and a General Research Fund (GRF)/Early Career Scheme (ECS) (24201919) from the Research Grants Council (RGC) in Hong Kong.

