## Supplementary Information for "Optical activation of TrkB receptors"

**Figure S1**

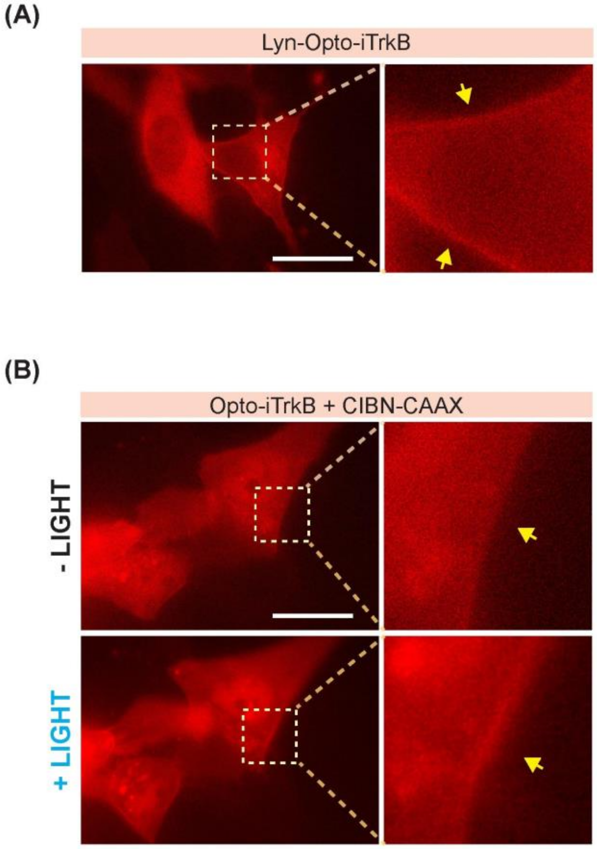


Fig. S1. (A) Lyn-Opto-iTrkB localized on the plasma membrane in NIH 3T3 cells expressing Lyn-iTrkB-mCh-CRY2. (B) Opto-iTrkB (CRY2-mCh-iTrkB) can be successfully recruited to the plasma membrane via CRY2-CIBN interaction upon blue light. NIH 3T3 cells were co-transfected with CRY2-mCh-iTrkB and CIBN-CAAX and subjected to one pulse of blue light exposure (200 ms pulse duration at 9.7 W/cm$^2$). Scale bars, 20 μm.

**Figure S2**

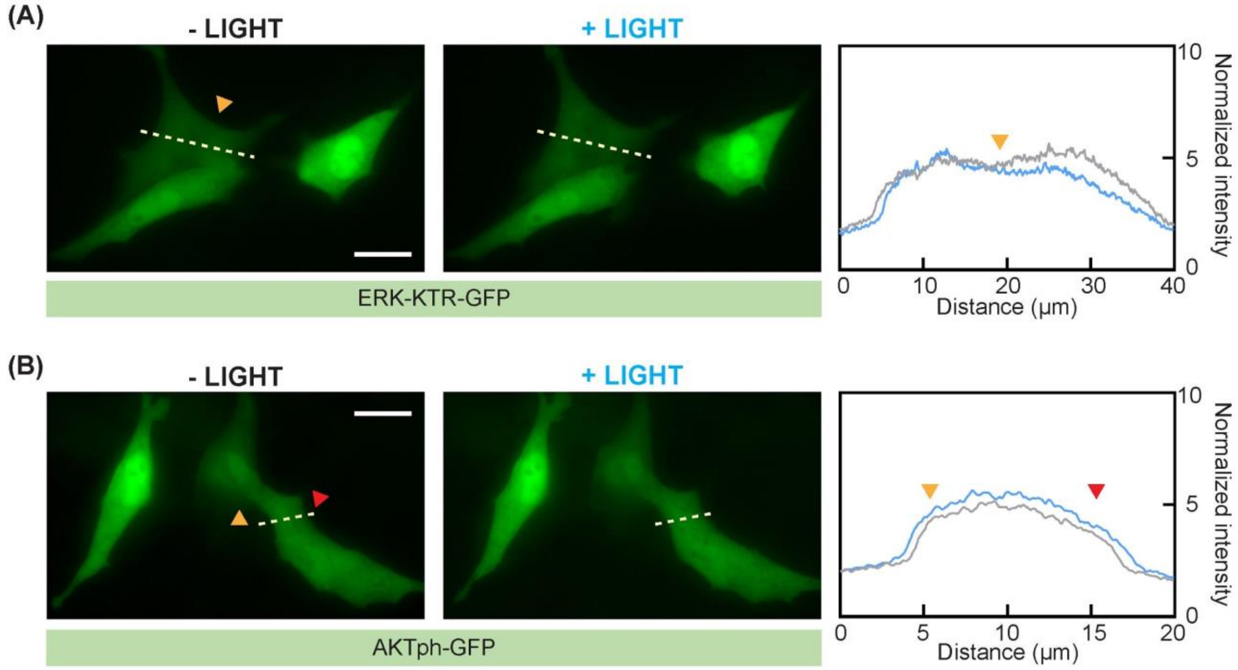


Fig. S2. No obvious translocation of ERK-KTR-GFP or AKTph-GFP was observed in cells without expressing OptoTrkB components. NIH 3T3 cells were co-transfected with (A) CIBN-CAAX and ERK-KTR-GFP; (B) CIBN-CAAX and Akt-GFP. Right panels show the normalized intensity change before (black line plot) and after blue light stimulation (blue light plot). Scale bars, 20 μm.

**Figure S3**

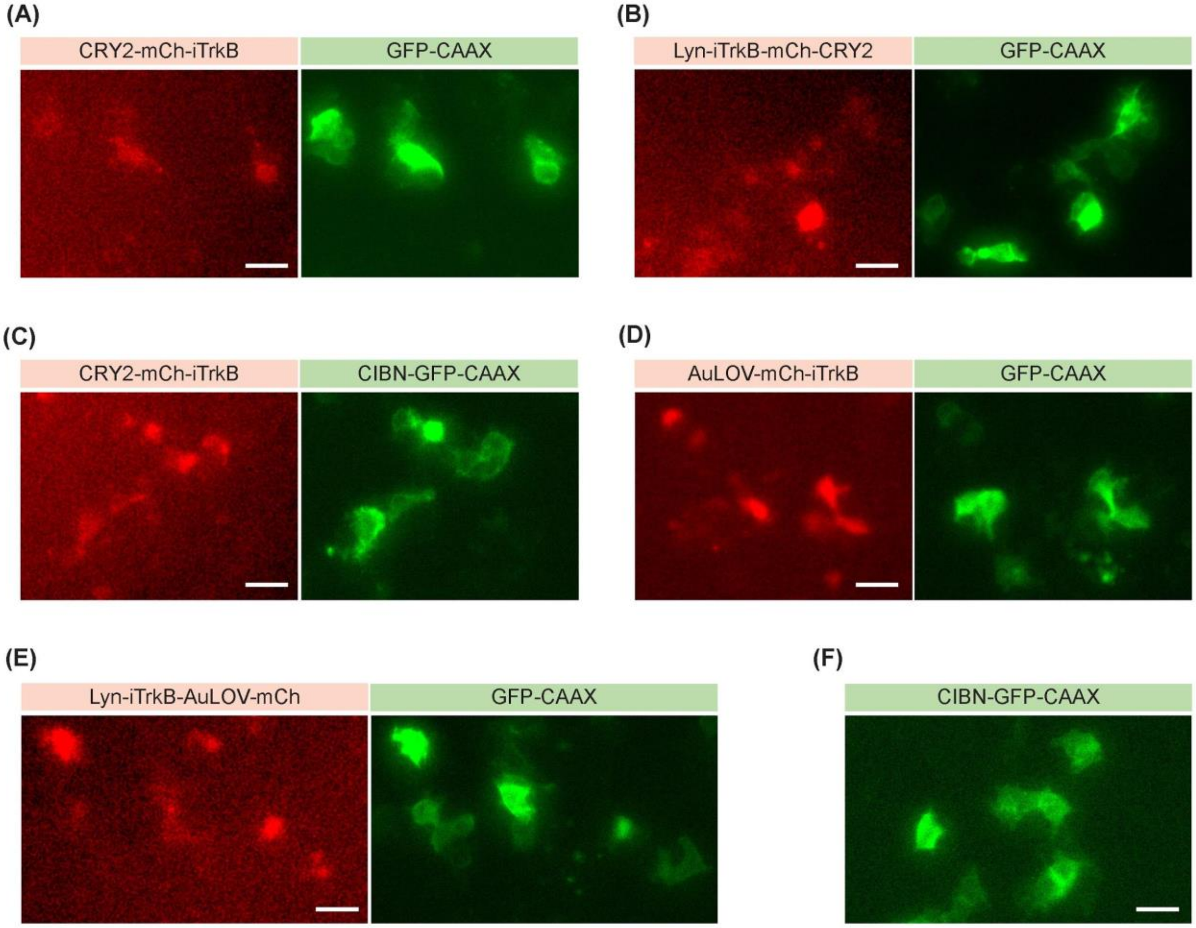


Fig. S3. PC12 cells expressing each of (A-C) CRY2-based OptoTrkB systems, (D-E) AuLOV-based OptoTrkB systems and (F) CIBN-GFP-CAAX showed no obvious neurite growth when kept in the dark. Scale bars, 20 μm.

**Figure S4**

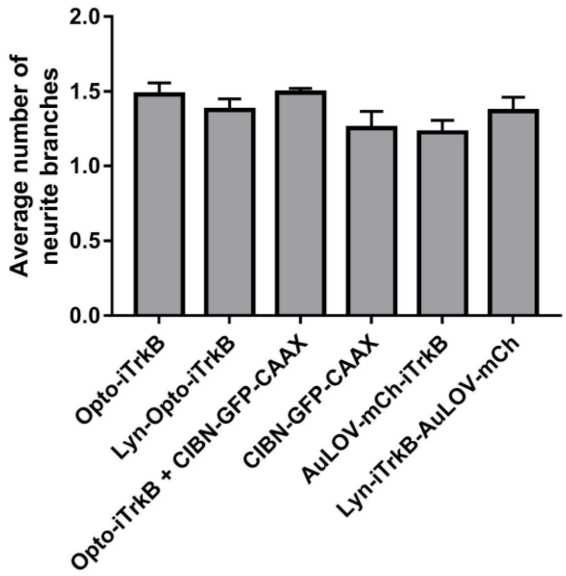


Fig. S4. Quantification of the average number of neurite branches in cells bearing neurites. Results are averaged from over 300 cells in group in three independent sets of experiments. Values represent mean ± SEM.

**Figure S5**

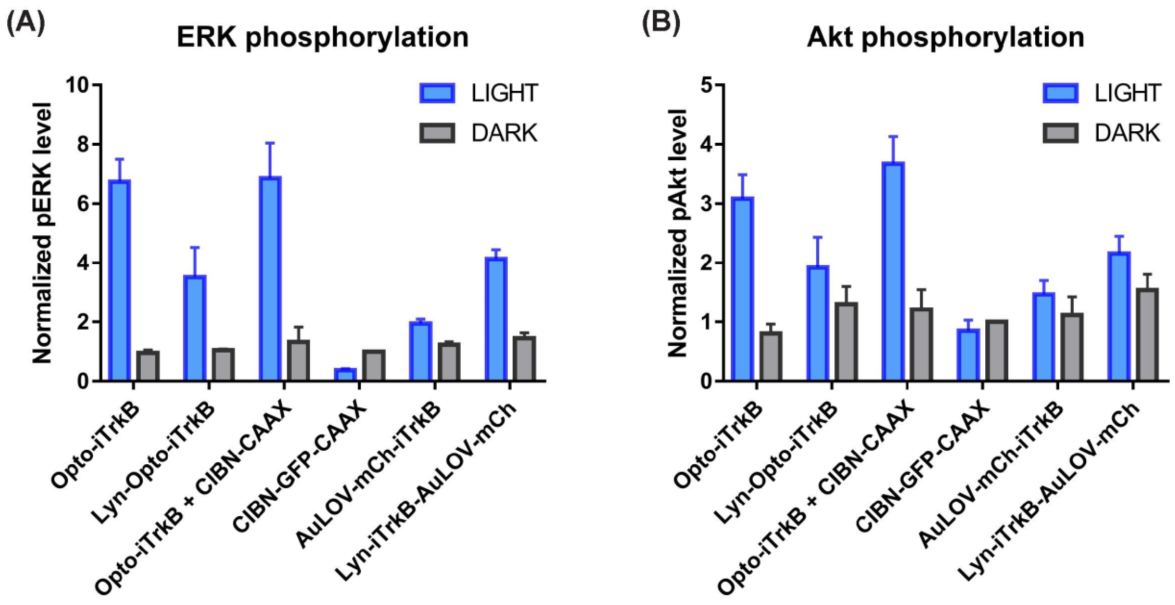


Fig. S5. Quantification of (A) ERK and (B) Akt phosphorylation level by densitometry analysis. pERK and pAkt levels were normalized to total ERK or total Akt level. Results were further normalized to the calculated pERK or pAkt level in the CIBN-GFP-CAAX control group kept in the dark. Plotted values represent the mean ± SEM of three technical replicates across two independent biological experiments.

**Table S1**

| Group | | Set 1 | | Set2 | | Set3 | |
|---|---|---|---|---|---|---|---|
| | | Total # | Neurite+ | Total # | Neurite+ | Total # | Neurite+ |
| CRY2-mCh-iTrkB | LIGHT | 332 | 49 | 317 | 67 | 430 | 64 |
| | DARK | 373 | 19 | 321 | 17 | 344 | 31 |
| Lyn-iTrkB-mCh-CRY2 | LIGHT | 366 | 45 | 335 | 63 | 399 | 53 |
| | DARK | 379 | 36 | 320 | 32 | 347 | 24 |
| CRY2-mCh-iTrkB + CIBN-GFP-CAAX | LIGHT | 363 | 123 | 357 | 124 | 342 | 152 |
| | DARK | 328 | 38 | 308 | 30 | 259 | 57 |
| AuLOV-mCh-iTrkB | LIGHT | 350 | 24 | 356 | 43 | 403 | 36 |
| | DARK | 544 | 38 | 311 | 17 | 414 | 32 |
| Lyn-iTrkB-AuLOV-mCh | LIGHT | 368 | 67 | 313 | 56 | 450 | 49 |
| | DARK | 376 | 34 | 351 | 28 | 327 | 25 |
| CIBN-GFP-CAAX | LIGHT | 400 | 27 | 308 | 18 | 579 | 55 |
| | DARK | 351 | 19 | 350 | 20 | 340 | 22 |

Table S1. Counting numbers for the quantification of PC12 cell differentiation. Cells bearing neurite growth >10 μm long were counted as cells with neurite growth (Neurite +) among total transfected cells (Total #).
